# Detailed investigation on the role of lipid metabolizing enzymes in the pathogenesis of retinopathy of prematurity among preterm infants

**DOI:** 10.1101/2022.05.13.491711

**Authors:** Saurabh Kumar, Satish Patnaik, Manjunath B Joshi, Subhadra Jalali, Komal Agarwal, Ramesh Kekunnaya, Subhabrata Chakrabarti, Inderjeet Kaur

## Abstract

**Purpose:** Extremely preterm infants are at risk of developing retinopathy of prematurity (ROP), characterized by an initial insufficient vascular network development in the retina (due to hyperoxia) that progress to neovascularization and neuroinflammation (hypoxic phase) ultimately leading to partial or total vision loss. Lipid metabolism has been shown to be a significant pathway that is involved in the regulation of angiogenesis, inflammation, and apoptosis in oxygen induced retinopathy mouse model, however, it is not explored in human ROP patients. The present study aimed to explore the association of lipid metabolizing, angiogenic and apoptotic genes with altered lipid metabolites in the ROP patients with different severity.

**Methods:** The blood, vitreous humor (VH), and fibrovascular membrane (FVM) samples were collected from premature infants diagnosed with ROP and controls. Gene expression of lipid metabolizing enzymes, angiogenesis, and apoptotic genes were performed using semi-quantitative PCR in blood. Lipid metabolites were identified and quantified by LC-MS in VH and were correlated with gene expression. The expression of key lipid metabolizing enzymes in severe stages of ROP was assessed by measuring their expression in FVM by immunohistochemistry.

**Results:** Genes coding for the lipid metabolizing enzymes such as *CYP1B1, CYP2C8, COX2*, and *ALOX15* were upregulated while *EPHX2* responsible for the conversion of epoxide fatty acids into diol fatty acids was significantly downregulated in ROP cases. The increase in the metabolic intermediates generated from the lipid metabolism pathway further confirmed the role of these enzymes in ROP. except for *EPHX2* which did not show any change in its activity. The glial cells in the FVM of ROP infants too showed a lack of EPHX2 expression. A significantly higher expression of genes involved in angiogenesis (*VEGF165/189, NOTCH1*, and *APH1B*) and apoptosis (*CASP3/8*) correlated with altered activity of lipid metabolizing enzymes (based on metabolites levels) among ROP cases.

**Conclusions:** Lipid metabolism may play a significant role in ROP development and progression. EPHX2 activity is a key step in the metabolic pathway of arachidonic acid that mediates and regulates inflammation and vascular pathology in preterm infants.

## INTRODUCTION

Retinopathy of prematurity (ROP) is a complication of preterm born infants with abnormal blood vascularization in the retina, that could eventually lead to fibrovascular retinal detachment. Such preterm infants are treated with laser and/or anti-VEGF therapy. Approximately 10% of preterm infants born annually are turned blind as a result of ROP in high-income countries while 40% of preterm infants are reported affected in low- and middle-income countries (1–3). The current treatment strategies come with adverse side effects and cannot prevent the reoccurrence of the disease. Moreover, these preterm ROP infants are also at a higher risk for developing certain eye problems later in life, such as retinal detachment, myopia (near-sightedness), strabismus (crossed eyes), amblyopia (lazy eye), and glaucoma. Another potential contributor to the pathology is a deficiency of ω-3 PUFAs, particularly DHA (3, 4). The macular layer in the retina is rich in long-chain polyunsaturated fatty acids (PUFAs) such as arachidonic acids (AA), Linoleic Acid (LA), docosahexaenoic acid (DHA), eicosapentaenoic acid (EPA), etc. which are prone to frequent lipid peroxidation and structural modification (5). These PUFAs and lipid metabolizing enzymes such as cyclooxygenases, lipoxygenases, and hydrolases have been studied widely for their role in cancer and cardiovascular diseases however, their involvement in ROP has been less explored. The activity of COXs and LOXs on arachidonic acid generate prostaglandins (PGs), thromboxane A2 (TXA2), leukotrienes (LTs), and hydroxy eicosatetraenoic acid (HETEs) which are the source of inflammation and hypertension (6, 7). Further, AA is also acted upon by a group of CYPs enzymes (ω-hydroxylases and epoxygenase pathway) generating HETEs and eicosatetraenoic acid (EETs) which regulate some cellular processes of carcinogenesis and progression, including cell proliferation, survival, angiogenesis, invasion, and metastasis. The EETs are mainly metabolized by soluble epoxide hydrolase (sEH) to the corresponding diols or dihydroxyeicosatrienoic acids (DHETs).

These diol fatty acids serve as important structural and signal mediating molecules in the endothelial layer, and immune cells, maintaining optimal retinal physiology and anatomy (8). Hu et al. reported that diols synthesized from PUFAs by the action of lipid metabolizing enzyme regulate angiogenesis by interfering Notch signaling pathway). Presenilin 1 and γ-secretase activity in oxygen-induced retinopathy model with sEH^-/-^ mutant mice is dysregulated which induces angiogenesis (9). Hu *et al* have reported the role of astrocyte apoptosis and muller glial cells being affected by a reduced activity of sEH in the retina of OIR models leading to neuroinflammation and neovascularization (9, 10). ROP is clinically diagnosed with symptoms of neuroinflammation, neovascularization, loss of retinal cells, and formation of the fibrovascular membrane (FVM). Any alteration at the level of expression of genes or metabolites can lead to irregular neovascularization of blood vessels, leading to severe ROP stages.

Hypoxia is extensively explored as a major factor in the development of ROP and its progression, however, the role of PUFA-related metabolites and enzymes that regulate angiogenesis has not been studied in human patients though few studies have looked at these in animal models. Our study is an attempt to systematically understand the role of arachidonic acid metabolism in ROP pathogenesis by integrated targeted transcriptomics and fatty acid metabolomics in premature infants. Solakivi *et al*. (11) had reported that PUFA such as arachidonic acid increases MMP9 secretion in human monocytic cell lines. A previous SNP-based genetic screening of ROP cases from our lab too demonstrated the significant association of genes involved in lipid metabolism, matrix metalloproteinase (MMP 9 and MMP2), and complement pathway activation in the progression of ROP. In this study, we focussed primarily on understanding the role of fatty acid metabolism, especially the activity and expression of the *EPHX2* gene that encodes for lipid metabolizing enzyme soluble epoxy hydrolase (sEH), in ROP pathogenesis. While there are a few studies on global metabolomics in plasma samples of ROP patients (12) and metabolomics of retina from oxygen-induced mice model of ROP (13), these do not reflect the actual complex metabolic profiles in the human eyes. Human vitreous humor has been extensively used as surrogate tissue being localized to the posterior retina and can reflect any damage to the retina based on the protein and metabolites secreted from retinal cells into the vitreous humor. Rathi *et al*. reported the presence of macrophages and neutrophils in the vitreous humor of ROP (14). These immune cells are also known to be involved in fatty acid metabolism. Continuing on our earlier findings on ROP genomics and proteomics, herein we propose to further understand the interplay of genes involved in lipid metabolism, inflammation, angiogenesis, and cell death (*CYP1B1, CYP2C8, COX2, ALOX15, EPHX2, VEGF165/189, PSEN1, APH1B, NOTCH1, CASP3/8*) from leukocytes of the preterm infants with and without ROP and correlated their expression with the metabolites in the vitreous humor. A functional evaluation of lower expression of *EPHX2* was assessed by immunostaining of the enzyme-soluble epoxide hydrolase in the fibrovascular membrane formed at the vitreoretinal junction of ROP infants. Our study establishes an important role for the enzymes involved in omega 3 fatty acid metabolism in the progression of ROP among preterm infants. This information can also serve as a major resource for developing better therapeutic options for ROP.

## MATERIAL AND METHODS

### Study Subjects

The study protocol adhered to the principles of the declaration of Helsinki and was approved by the Institutional Review Board of the L V Prasad Eye Institute (LVPEI), Hyderabad. Preterm babies referred for further management from the neonatal intensive care units of different hospitals to the LVPEI were enrolled in the study. The study cohort for serum samples comprised 126 preterm infants of GA ≤ 35 weeks and/or BW ≤ 1,700 g with severe ROP (*n* = 70) and control or moderate ROP (*n* = 56). The study cohort for vitreous humor samples comprised 15 preterm babies with GA ≤ 35 weeks and/or BW ≤ 1,700 g with severe ROP and 15 full-term mature infants with GA ≥ 35 weeks and /or BW ≥ 1500 with congenital cataract. The detailed demographic and clinical history of all the infants recruited for the study was documented and written informed consents were obtained from their guardians. The diagnosis and categorization of ROP cases were done based on severity (stages 1–5), location (zones I, II, III), amount of disease (clock hours), and presence or absence of “plus” disease following ICROP guidelines. Severe ROP includes progressive disease and highly vascularized retina, which requires prompt treatment. No/Mild ROP cases include less severe disease, which does not require any treatment and have no neo-vascularization compared to severe ROP patients. The control infants and the infants with mild ROP were considered to have no neo-vascularization and are considered control while infants with stage 4 and stage 5 with plus who have high neo-vascularization are considered infants with severe ROP in this study. Although until the regression of the disease completely, infants were under regular follow-up for ROP screening.

### Sample Collection

Venous blood (0.5–1.5 mL) was collected from the preterm infants diagnosed with severe ROP and no/mild ROP by venipuncture. RNA was extracted from the blood samples using Trizol and an automated cDNA extraction platform following the manufacturer’s guidelines. Likewise, for metabolomics studies, the vitreous humor samples (400–500 μL) were collected from preterm infants with stage IV and V ROP (*n* = 15) who had undergone vitrectomy as a part of their routine clinical management. The controls for the metabolomics studies included infants with congenital cataracts (<6 months of age) who underwent partial vitrectomy as part of the surgical management (*n* = 15). Additionally, FVM were collected from preterm infants with stage IV and V ROP who had undergone membranectomy as a part of their routine clinical management. Human skin tissue was used as a positive control against the EPHX2 antibody.

### Materials

All the primers for semi-quantitative PCR were purchased from Bioserve Biotechnologies (India) Pvt Ltd. The rabbit polyclonal antibody against soluble epoxide hydrolase (sEH) was purchased from R&D Biosystems Inc., USA and the mouse monoclonal antibody anti-Glial fibrillary acidic protein (GFAP) was purchased from Cell Signaling Technology, USA for immunohistochemistry (IHC). Secondary antibodies for IHC were from LI-Cor Biosciences. All other chemicals (unless otherwise specified) were from Sigma-Aldrich.

### Semi-Quantitative PCR

RNA from serum samples of severe ROP and no/mild ROP control was extracted by the QIAamp RNA Blood Mini kit method (Qiagen, Hilden, Germany). The cDNA was prepared using the iScript cDNA synthesis kit (Bio-Rad, CA, USA). Semi-quantitative PCR was carried out using the specific primers (Supplementary Table 1) for *CYP1B1, CYP2C8, COX2, ALOX15, EPHX2, VEGF165/189, PSEN1, APH1B, NOTCH1, and CASP3/8* while *β-actin* was used as an endogenous control.

### FVM preparation and analysis

To determine the expression of EPHX2 and GFAP in ROP eyes, fibrovascular membranes (FVM) from preterm infants, retinal control tissue, and human skin tissue were immersed in 4% formalin for 24 hours and embedded in a paraffin block and stored until use. The tissue section from the paraffin block was placed on a pathological glass slide and was deparaffinized. The antigen retrieval was performed using ammonium acetate buffer followed by brief heating. For whole-mount immunohistochemistry, FVM and skin tissue was fixed in 4% paraformaldehyde (PFA) for 20 minutes at room temperature and washed with PBS buffer thrice. After isolation, tissues were blocked and permeabilized in 2% BSA and 0.5% Triton X-100 at room temperature for 1 hour. The primary antibodies for sEH (1:100) and GFAP (1:500), were diluted in 2% BSA and kept at 4°C overnight. Afterward, Alexa Fluor–coupled secondary antibodies (1:300) were used. Cell nuclei were visualized with DAPI (1:200; Molecular Probes, Thermo Fisher Scientific USA). All high-resolution fluorescent images were captured.

### Metabolomics analysis

#### Sample Preparation and Processing

Vitreous humor samples from Severe ROP (Stage IV/V) and control (congenital cataract) were collected at the time of vitrectomy from infants. Samples were collected in fresh vials and immediately stored in ice (Note-All samples were collected between 9 AM to 1 PM) and were proceeded for further processing within an hour of collection. Samples were centrifuged for 5 minutes at 14,000 g at 4°C and supernatant was carefully transferred into fresh cryotubes. 100 μl of vitreous humor was taken from the collected supernatant and 400 μl of methanol was added to it, mixed well, incubated in ice for 5 minutes, snap-frozen using liquid nitrogen, and then stored at −80C for further use. Small molecules were extracted using the cold methanol method as described earlier (Patnaik et al, 2019). Briefly, lyophilized samples were mixed with 200μl methanol and incubated in ice for 20 minutes, and centrifuged for 10 minutes at 4°C, at 13000 RPM. The supernatant was collected, lyophilized and 30 ul of acetonitrile containing 0.1% formic acid was added and subjected to mass spectrometry. The samples were run in Agilent 6520 ESI/QTOF mass spectrometry (Agilent 6520, Santa Clara, USA) coupled to an Agilent 1200 liquid chromatography system. Metabolites were separated using a reverse phase C18 column and analyzed over a range of 40–1200 m/z. Basic and neutral metabolites were eluted in positive mode at 400 μL/min flow rate using mobile phase A: 0.1% formic acid in water and mobile phase B: 0.1% formic acid in 90% acetonitrile (2% to 98% B in 25 min, 98% B for10 min and equilibrated to 2% B for 10 min). The parameters applied for mass detection included: nitrogen gas flow rate-8 l/min, gas temperature-330° C, nebulizer gas pressure-35 psi, Vcap-3700 V, fragmentor-160V, skimmer-65V, mass scan range-m/z 50–1000.

#### Metabolomics data processing

Spectral features were obtained from raw data using molecular feature extraction (Agilent, Santa Clara) and were aligned using XCMS. We annotated the small molecules Human Metabolome Data Bank (HMDB) considering 15 ppm differences.

### Statistical analysis

Lipid-related metabolites were categorized into inflammatory and anti-inflammatory-derived metabolites. Fold change and P-value were calculated for the same using Students’ T-tests. Also, performed predictive analysis for *EPHX2* gene and epoxy derived metabolite (LC-MS data) interactions using the MetScape-Cytoscape analysis tool.

## Results

### Study Cohort

126 preterm infants were recruited and analyzed for gene expression profiling. Their birth and clinical description are provided in Table 1 and Figure 1a. Quite expectedly, infants with ROP had a significantly lower gestational age and birth weight as compared to those with no ROP. The study population for metabolomics study from vitreous humor samples included 30 infants (ROP: 15; controls: 15). For a better understanding of the results, we have described these in three separate sections as follows:

> *Section A: Differential expression of genes involved in lipid metabolism, angiogenesis, inflammation, and cell death for understanding the role of lipid metabolizing enzymes in ROP pathogenesis*

**Table 1:**
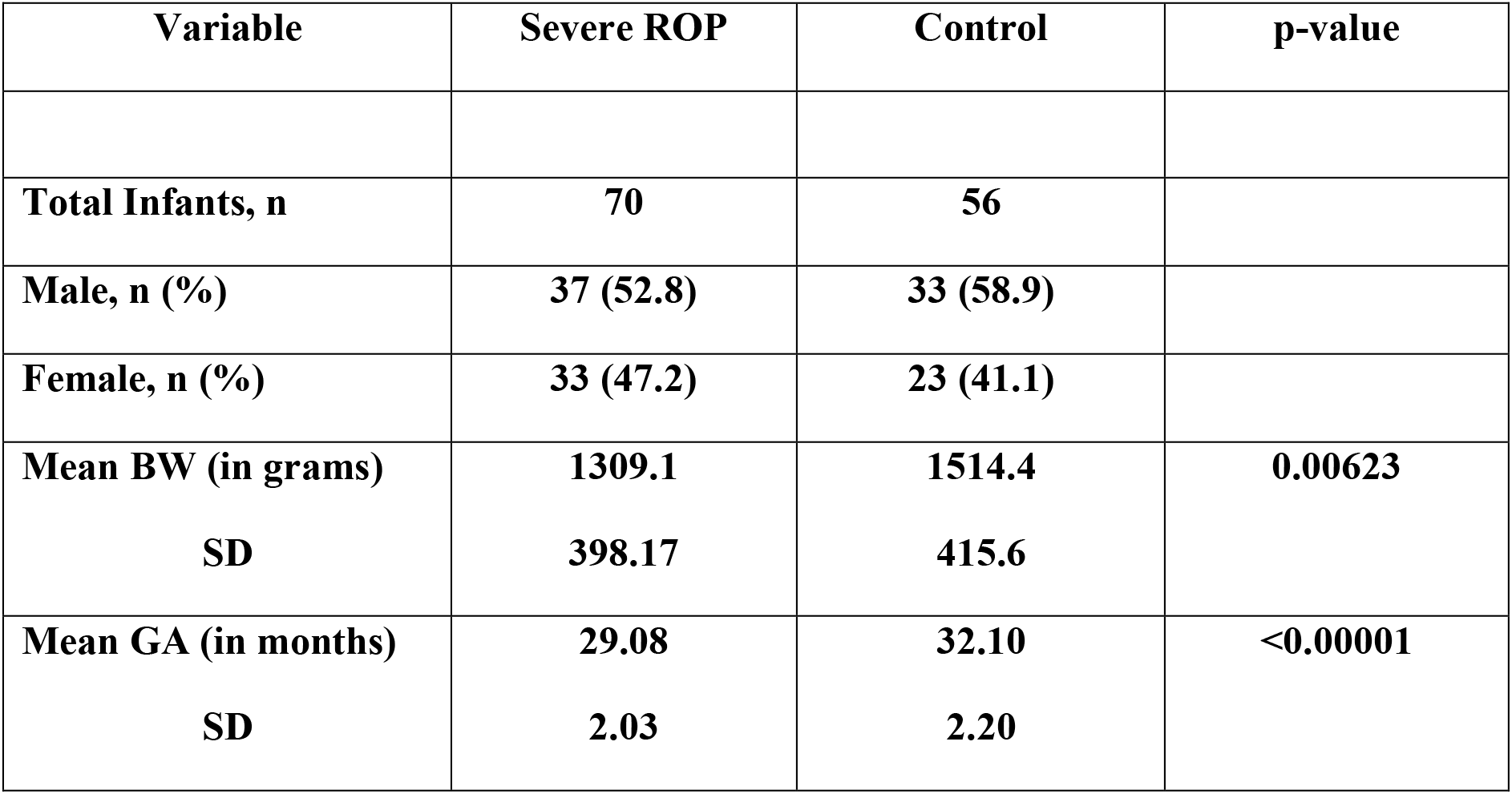
Details of the patients and controls recruited for the gene expression profiling in blood. (BW=birth weight; GA= Gestational Age: ROP = retinopathy of prematurity; SD = standard deviation.)

| Variable | Severe ROP | Control | p-value |
| --- | --- | --- | --- |
| **Total Infants, n** | **70** | **56** |  |
| **Male, n (%)** | **37 (52.8)** | **33 (58.9)** |  |
| **Female, n (%)** | **33 (47.2)** | **23 (41.1)** |  |
| **Mean BW (in grams)** | **1309.1** | **1514.4** | **0.00623** |
| **SD** | **398.17** | **415.6** |  |
| **Mean GA (in months)** | **29.08** | **32.10** | **<0.00001** |
| **SD** | **2.03** | **2.20** |  |

**Table 2:** Details of the patients and controls recruited for the metabolite profiling in the vitreous humor. (BW=birth weight; GA= Gestational Age: ROP = retinopathy of prematurity; SD = standard deviation.)

| Variable | Severe ROP | Control |
| --- | --- | --- |
| **Total Infants, n** | **15** | **15** |
| **Male, n (%)** | **7 (46.7)** | **9 (60)** |
| **Female, n (%)** | **8 (53.3)** | **6 (40)** |
| **Mean BW (in grams)** | **1286.9** | **2444.0** |
| **SD** | **237.3** | **531.5** |
| **Mean GA (in months)** | **29.8** | **37.5** |
| **SD** | **2.99** | **2.0** |

**Figure 1:**
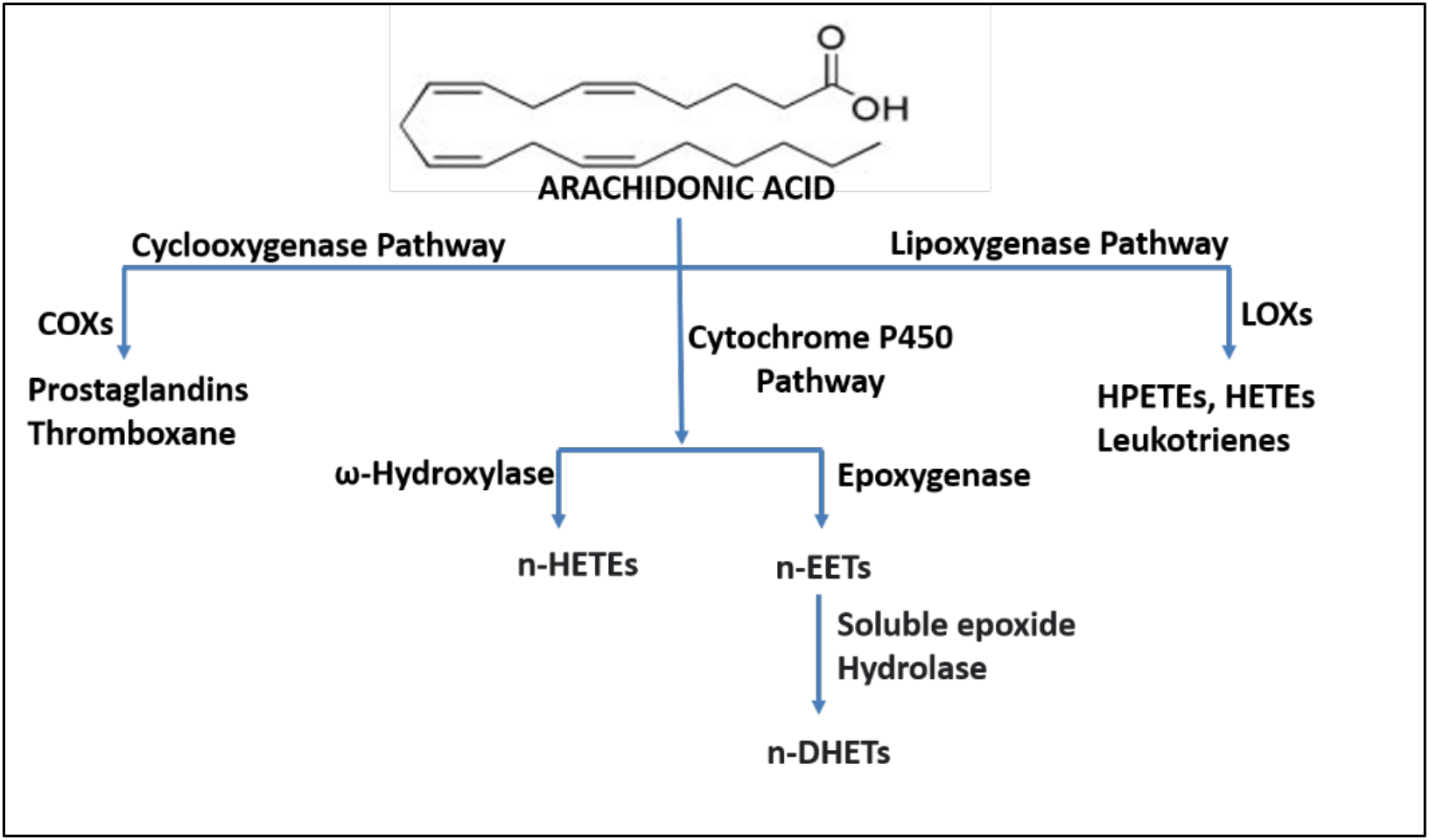
Arachidonic acid metabolism: Classification of different metabolites formed from arachidonic acid by the metabolism of the different enzymatic pathways. Cyclooxygenase and lipoxygenase pathway is mediated by the activity of COXs and LOXs respectively. Epoxygenase and ω-hydroxylase pathways are mediated by the activity of CYPs. EPHX2 (Soluble epoxide hydrolase) acts on the epoxygenase-generated metabolites to form dihydroxy alcohols.

#### Gene Expression of lipid metabolizing genes

The gene expression of lipid metabolizing genes, *COX2, ALOX15, CYP1B1, CYP2C8* and *EPHX2* was performed in serum of severe ROP and control samples (figure 2). Among these, prominent genes/enzymes involved in omega 3 fatty acid metabolism including *COX2, ALOX15, CYP1B1* and *CYP2C8* were significantly upregulated (3.85-fold, p-value= 0.05; 6.73-fold, p-value= 0.03; 4.9-fold, p-value= 0.007, 3.1-fold, p-value= 0.05 respectively) while *EPHX2* was significantly down regulated (1.57-fold, p-value= 0.04).

**Figure 2:**
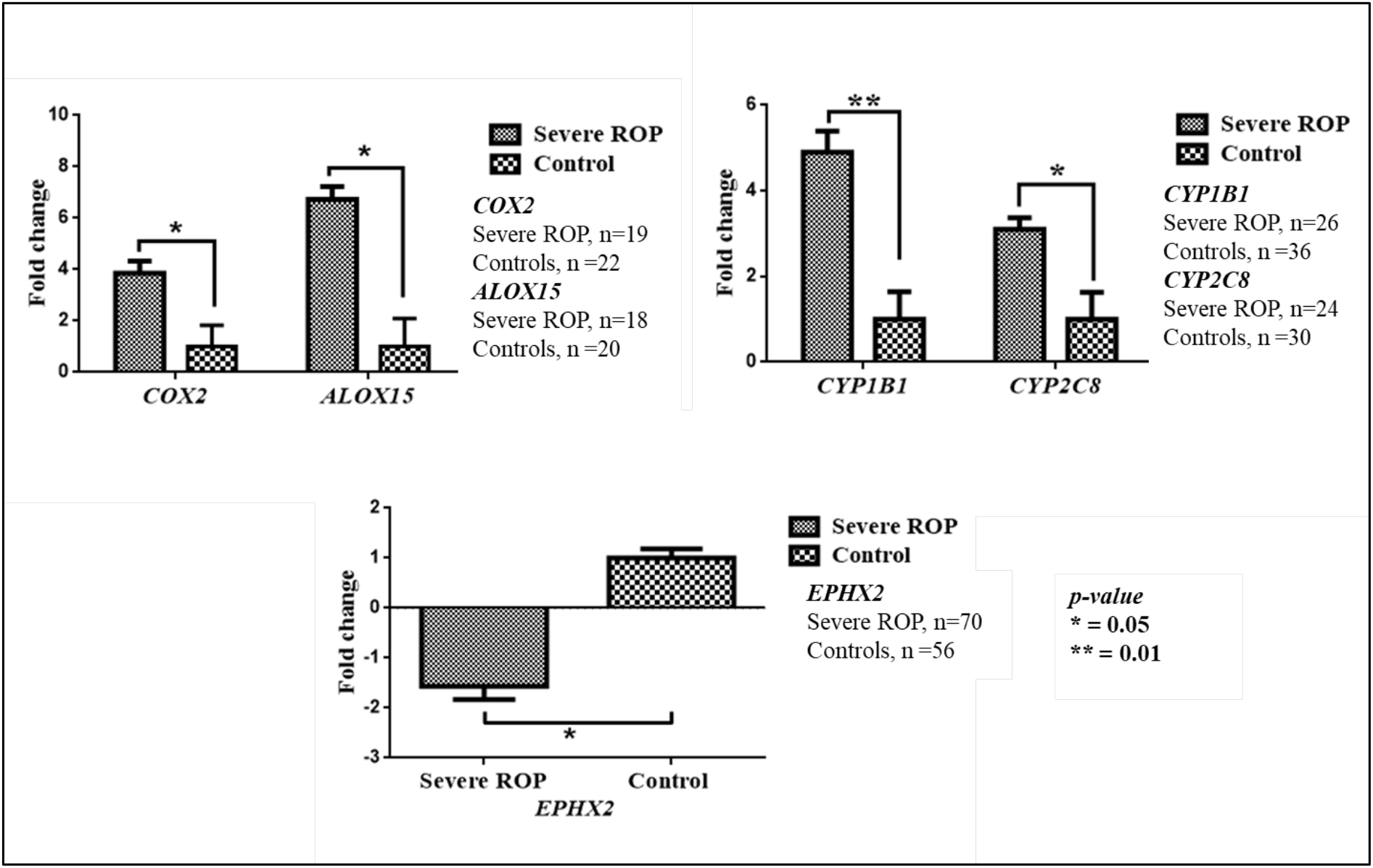
mRNA expression levels of lipid metabolizing genes. (A) qRT-PCR analysis of transcripts for COXs (severe ROP=19, control=22) and LOXs (severe ROP=18, control=20) expression (B) qRT-PCR analysis of transcripts for CYP1B1 (severe ROP=26, control=36) and CYP2C8 (severe ROP=24, control=30) expression (C) qRT-PCR analysis of transcripts for EPHX2 (severe ROP=70, control=50). All the graphs are representatives of biological triplicates. All graphs were prepared using GraphPad Prism version 8.0.2. β-actin was used as the normalization control for qRT-PCR. Statistical significance is represented as * and ** for p<0.05 and p<0.01 respectively.

#### Gene Expression of angiogenic genes

The gene expression of angiogenic genes, *VEGF165, VEGF189, PSEN1, APH1B* and *Notch1* was performed in serum of severe ROP and control samples (figure 3). As expected angiogenic genes including *VEGF165, VEGF189* and *APH1B* were significantly upregulated (7.24-fold, p-value= 0.001; 2.75-fold, p-value= 0.02; 4.86-fold, p-value= 0.05 respectively) while *PSEN1* was significantly (1.27-fold, p-value= 0.04) down regulated *Notch1* the regulator of angiogenesis was also significantly (6.6-fold, p-value= 0.003) upregulated.

**Figure 3:**
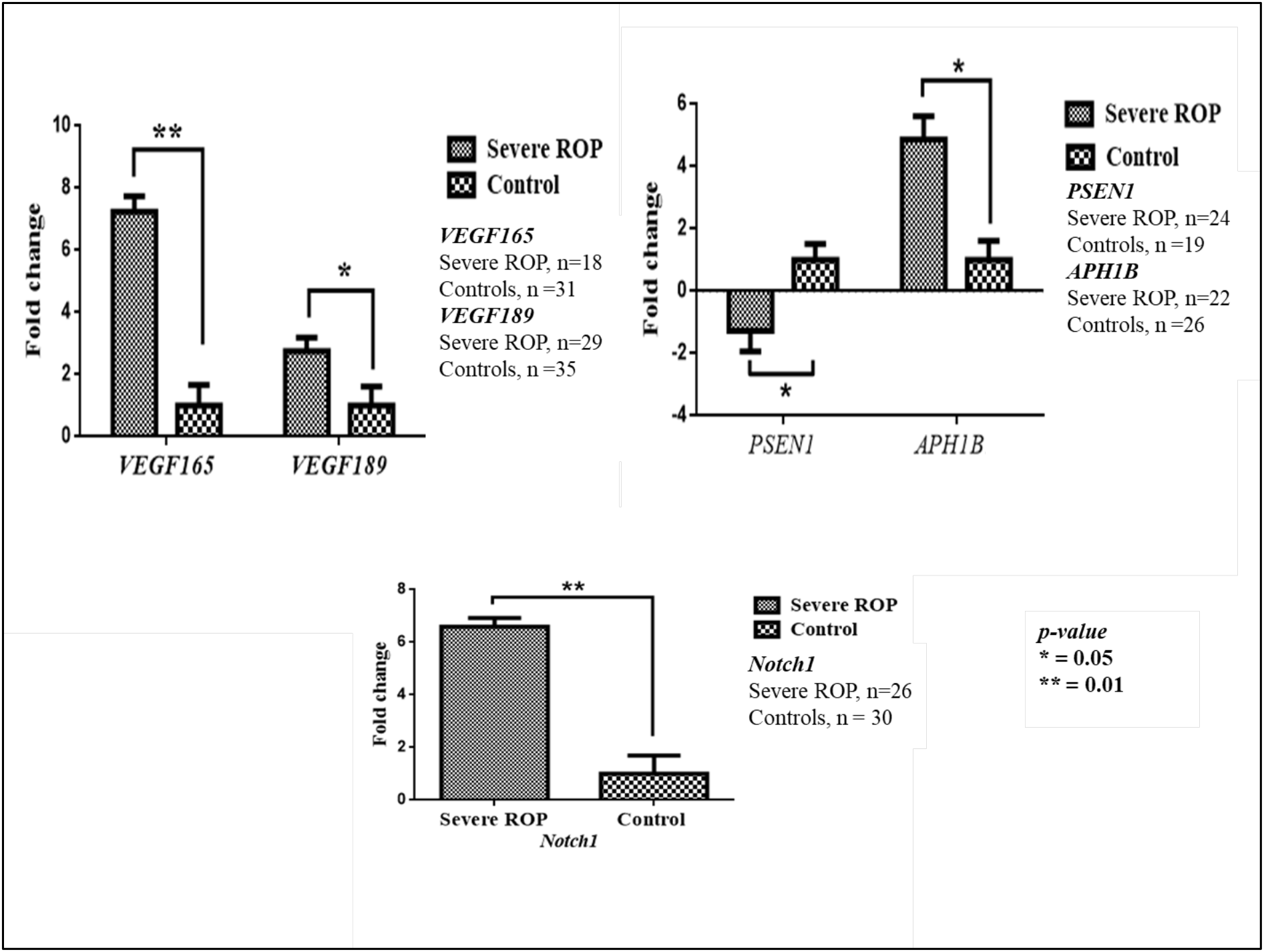
mRNA expression levels of angiogenic genes. (A) qRT-PCR analysis of transcripts for VEGF165 (severe ROP=18, control=31) and VEGF189 (severe ROP=29, control=35) expression (B) qRT-PCR analysis of transcripts for PSEN1 (severe ROP=24, control=19) and APH1B (severe ROP=22, control=26) expression (C) qRT-PCR analysis of transcripts for NOTCH1 (severe ROP=26, control=30). All the graphs are representatives of biological triplicates. All graphs were prepared using GraphPad Prism version 8.0.2. β-actin was used as the normalization control for qRT-PCR. Statistical significance is represented as * and ** for p<0.05 and p<0.01 respectively.

#### Gene Expression of apoptotic genes

The gene expression of apoptotic genes, *CASP3* and *CASP 8* were performed in serum of severe ROP and control samples (figure 4). Both cleaved fragments of *CASP3* and *CASP8* were significantly upregulated (3.82-fold, p-value= 0.01- and 3.86-fold, p-value= 0.004).

> *Section B: Metabolite profiling for assessing the activity of lipid metabolizing enzymes in ROP pathogenesis*

**Figure 4:**
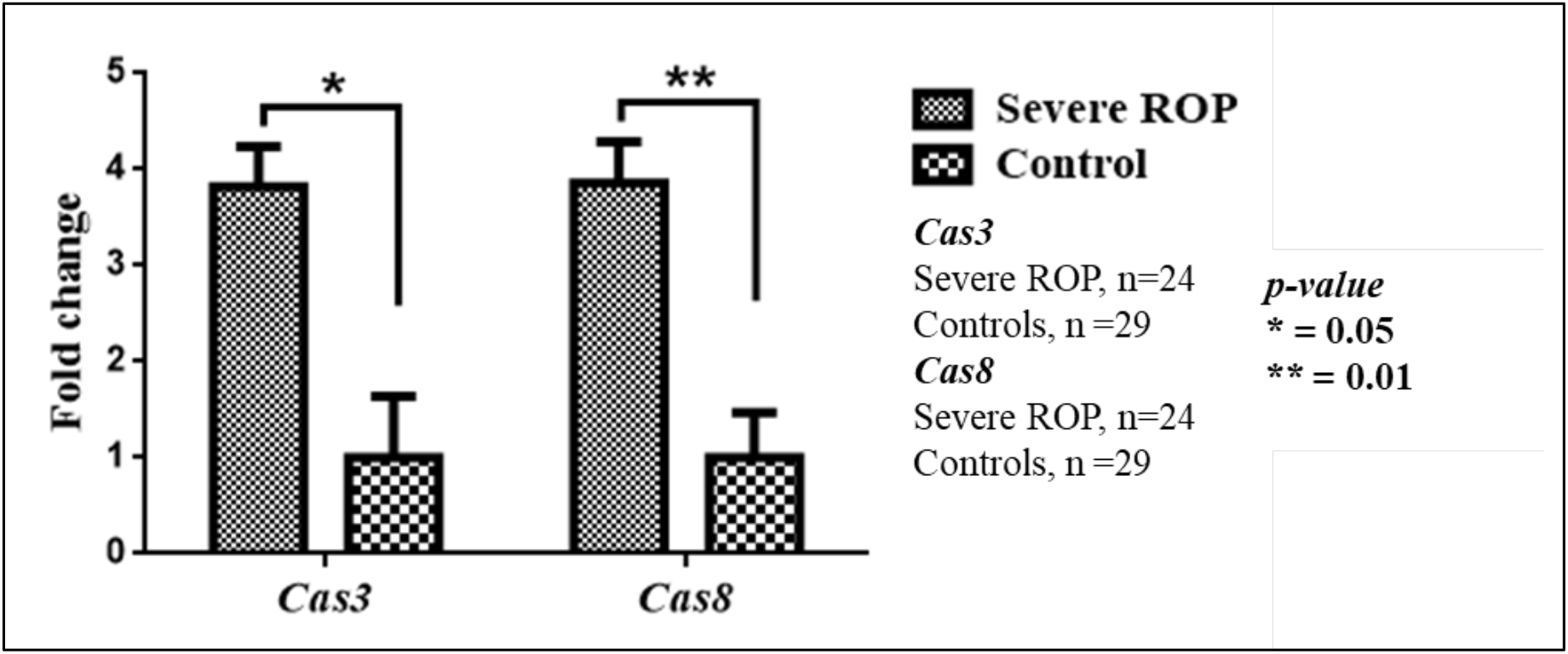
mRNA expression levels of apoptotic genes. (A) qRT-PCR analysis of transcripts for *CASP3* (severe ROP=24, control=29) and *CASP8* (severe ROP=24, control=29) expression All the graphs are representatives of biological triplicates. All graphs were prepared using GraphPad Prism version 8.0.2. β-actin was used as the normalization control for qRT-PCR. Statistical significance is represented as * and ** for p<0.05 and p<0.01 respectively.

#### LC-MS/MS for Metabolite Analysis

Inflammation-mediated metabolites (M341T34, M351T30, M353T23, M331T21, M366T31, M196T4, M287T33) were found to be significantly upregulated while anti-inflammatory metabolites were downregulated (Metlin ID M339T27) (Figure 5, Table 3).

**Figure 5:**
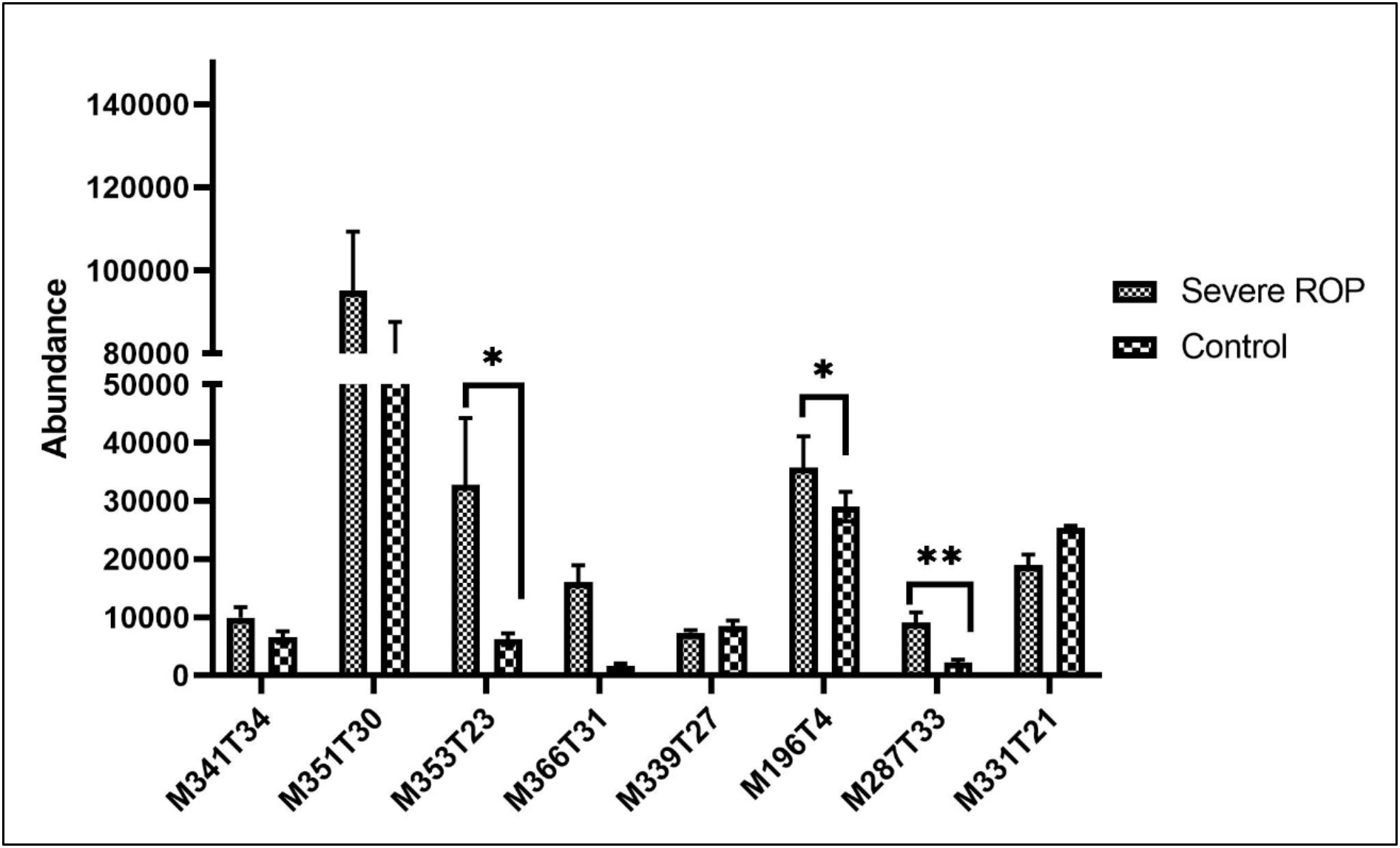
Abundance of fatty acid metabolites in vitreous humor. (A) The graphical representation of the abundance of metabolites detected in the vitreous humor of severe ROP (n=15) compared to control (n=15). The corresponding x-axis represents the Metlin ID for the metabolites detected. All the graphs are representatives of biological duplicates. All graphs were prepared using GraphPad Prism version 8.0.2. Statistical significance is represented as * and ** for p<0.05 and p<0.01 respectively.

**Table 3:**
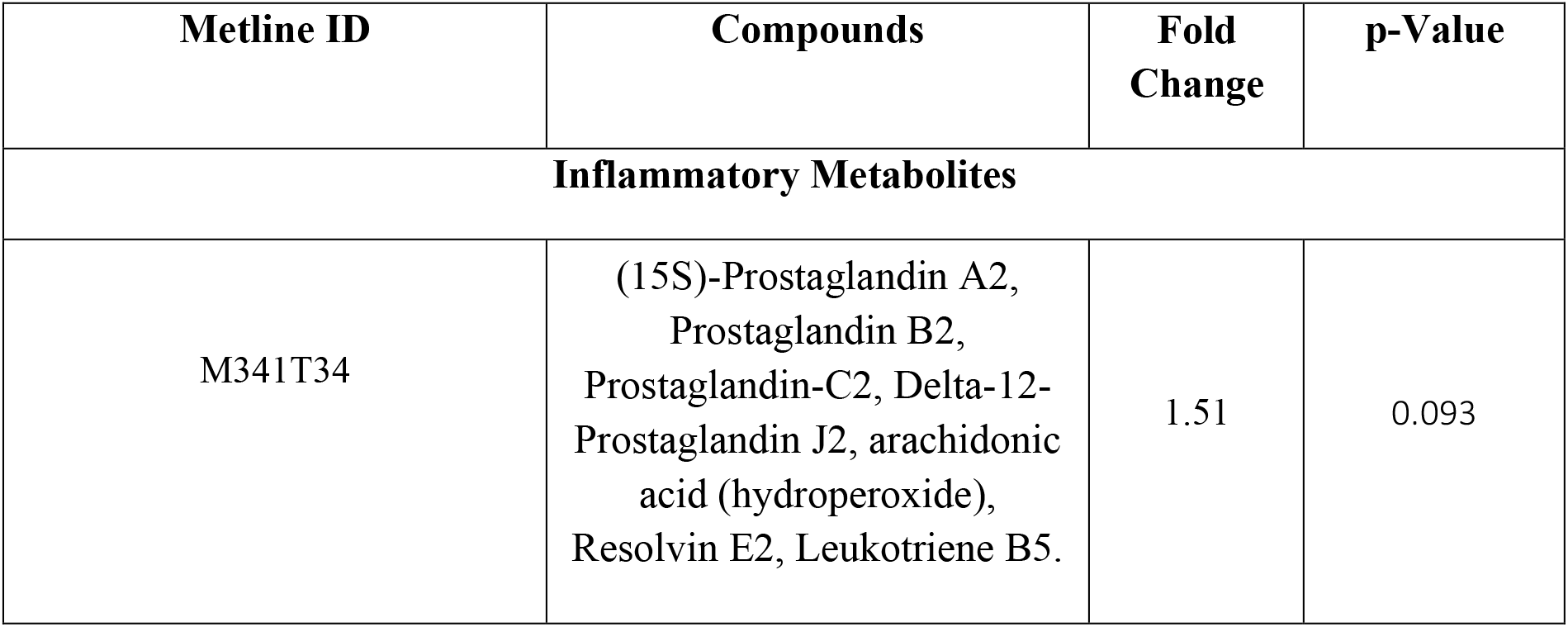
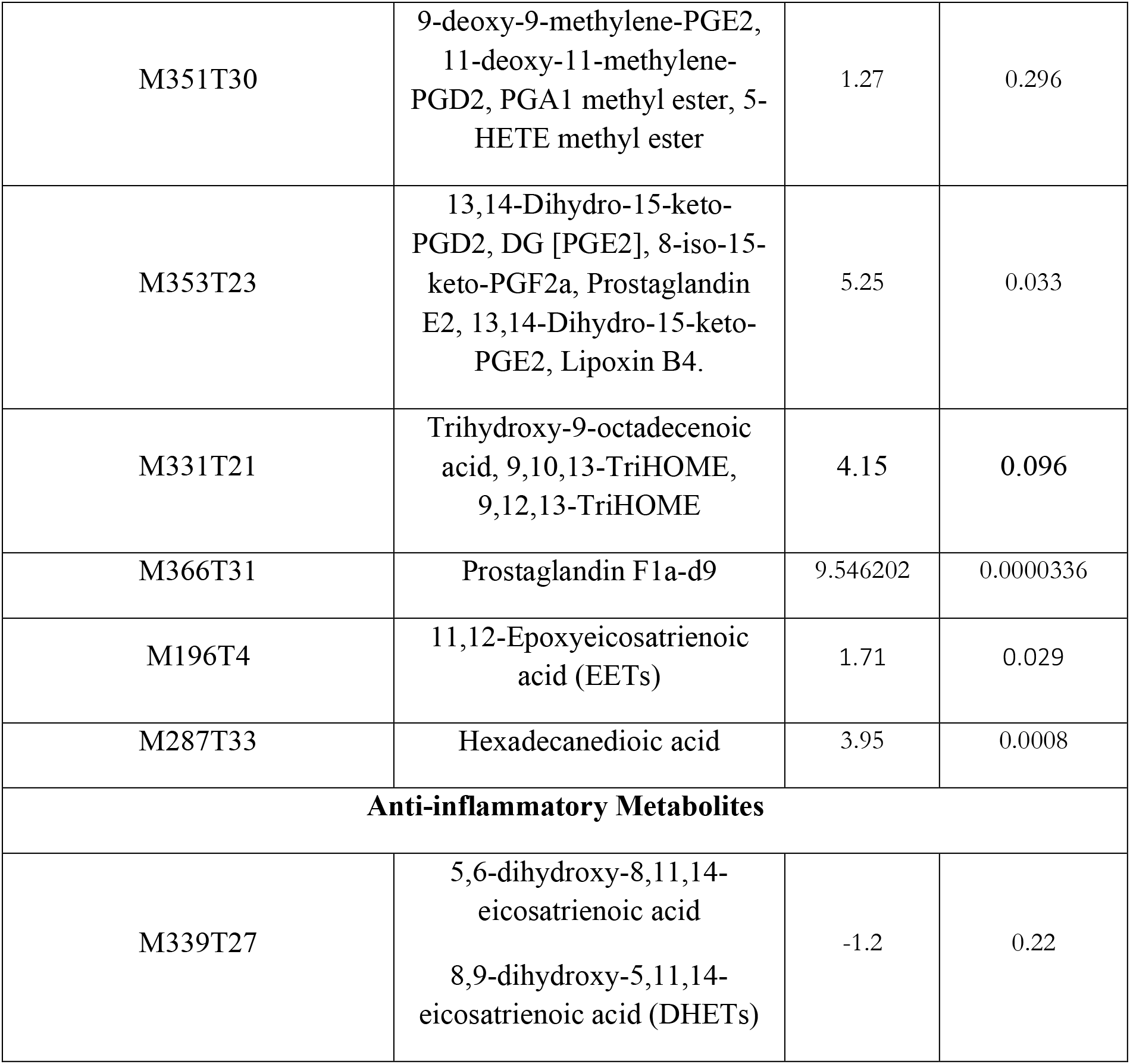
List of metabolites and their corresponding Metlin IDs observed in against peak is observed in the vitreous humor. The metabolites are further classified into inflammatory and anti-inflammatory metabolites. 15 ppm error was used for the analysis. P-value and fold change are denoted in the respective column.

#### Combined analysis of *EPHX2* Gene and metabolic interactions

Next to delineate the underlying mechanisms involving the *EPHX2* gene in the pathogenesis of ROP, an interactive map for the EPHX2 gene and epoxy-derived metabolites were constructed by using Metscape (Cytoscape plugin). *EPHX2* gene has both direct and indirect interactions with several inflammatory mediated metabolites and CYP genes (*CYP1B1, CYP2C8*, etc..). A majority of these gene and metabolites interactions have been shown involved in the different lipid metabolic pathways namely: arachidonic acid metabolism, Di-unsaturated fatty acid beta-oxidation, leukotriene metabolism, linoleate metabolism, putative anti-Inflammatory metabolites formation from EPA, and xenobiotics metabolism pathways (figure 6).

> *Section C: Further validation of EPHX2 activity in retinal tissues and in vitro model (primary retinal cells under hypoxia*)

**Figure 6:**
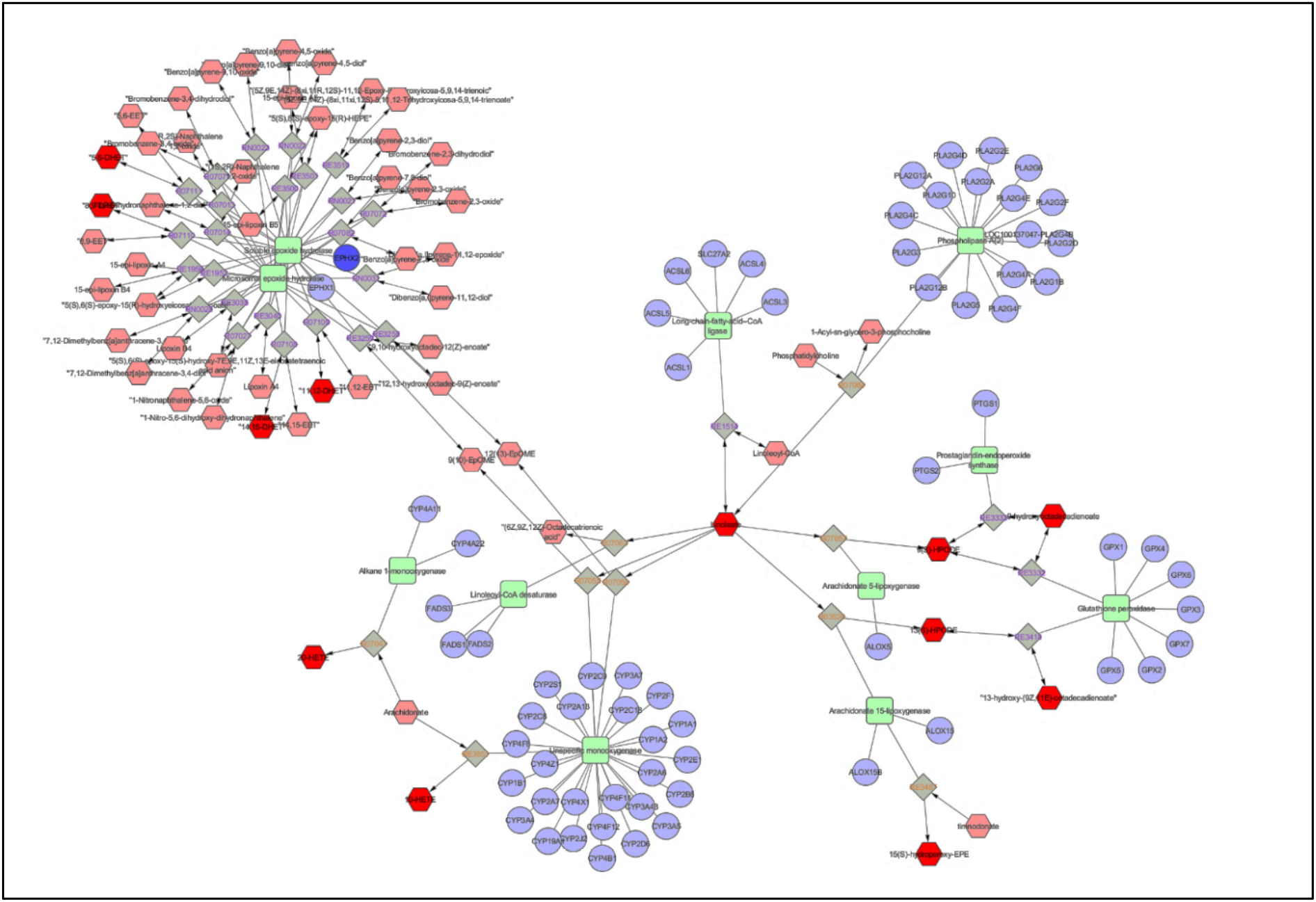
Network mapping of EPHX2 and lipid-derived metabolites. EPHX2 gene showing strong association with CYPs and other lipid metabolizing enzymes. The network map shows the synthesis reaction of different metabolites and their associations with the genes mapped.

**Figure 7:**
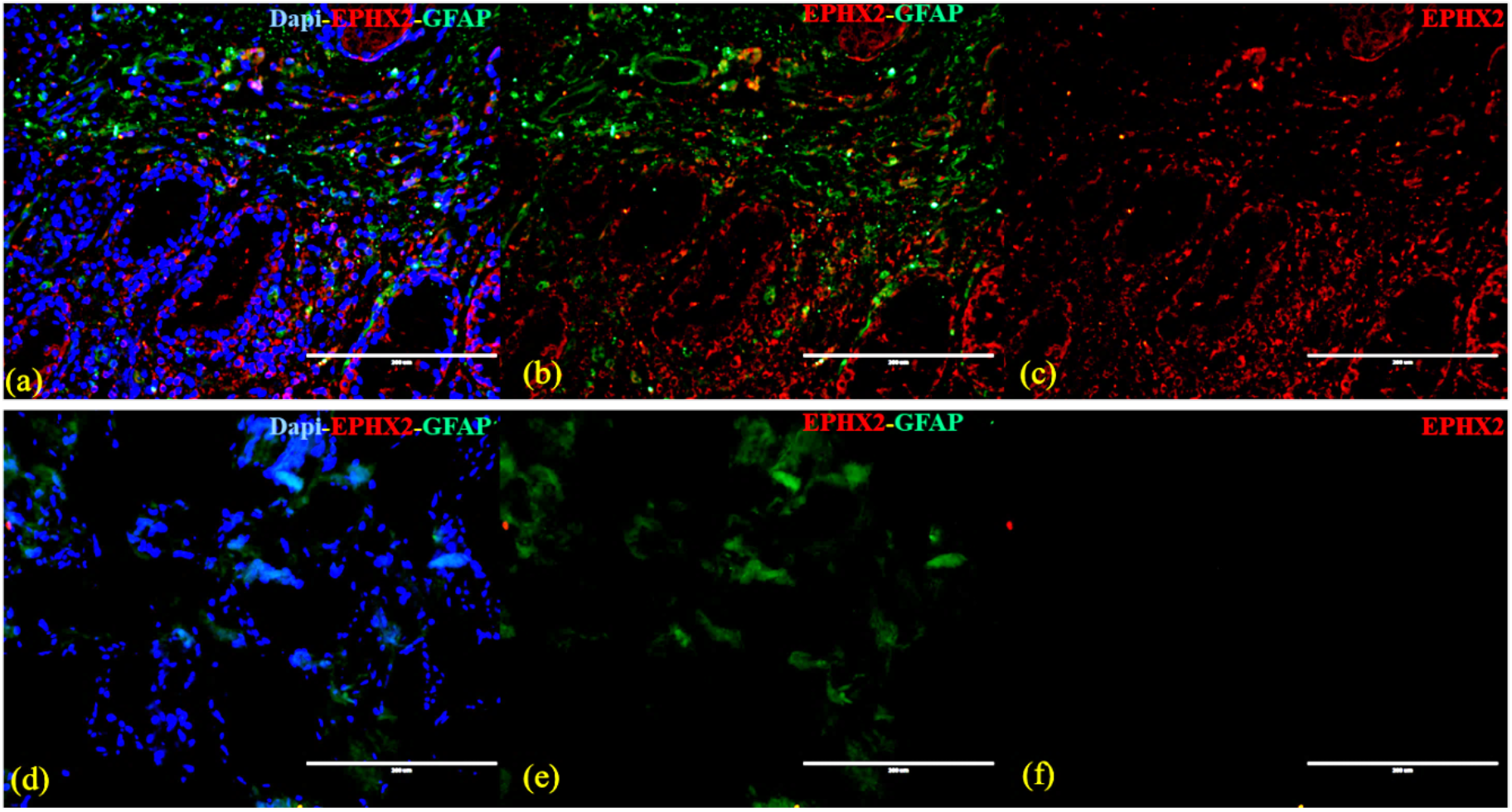
Immunohistochemistry of fibrovascular membrane and control. (A) Merge image of positive control showing expression of EPHX2 and GFAP when stained with DAPI. (B) Merge image of EPHX2 and GFAP showing expression in control. (C) EPHX2 expression in control. (D) Merge image of fibrovascular membrane showing expression of GFAP when stained with DAPI and no EPHX2 expression. (E) Merge image of EPHX2 and GFAP showing expression in FVM, but no expression of EPHX2 (F) FVM showing no expression of EPHX2. All the images are representatives of biological triplicates. Skin tissue was used as positive control and no antibody was used as a negative control.

#### Immunohistochemistry of FVM and human retina

Immunohistochemistry was performed for confirming the expression of EPHX2 protein in the astrocytes and other glial cells, RPE cells, and extension of retinal ganglion cells (FVM) from ROP eyes. The human retina and skin tissue are used as the positive control. Positive staining of astrocyte marker GFAP was seen in FVM in the different layers and the lining of the retinal blood vessel while no expression of EPHX2 was seen. The normal human retina showed both expressions of EPHX2 and GFAP in the cells lining the retinal blood vessels. The human skin tissue used as positive control showed high expression of EPHX2 (Figure 8a-g).

**Figure 8:**
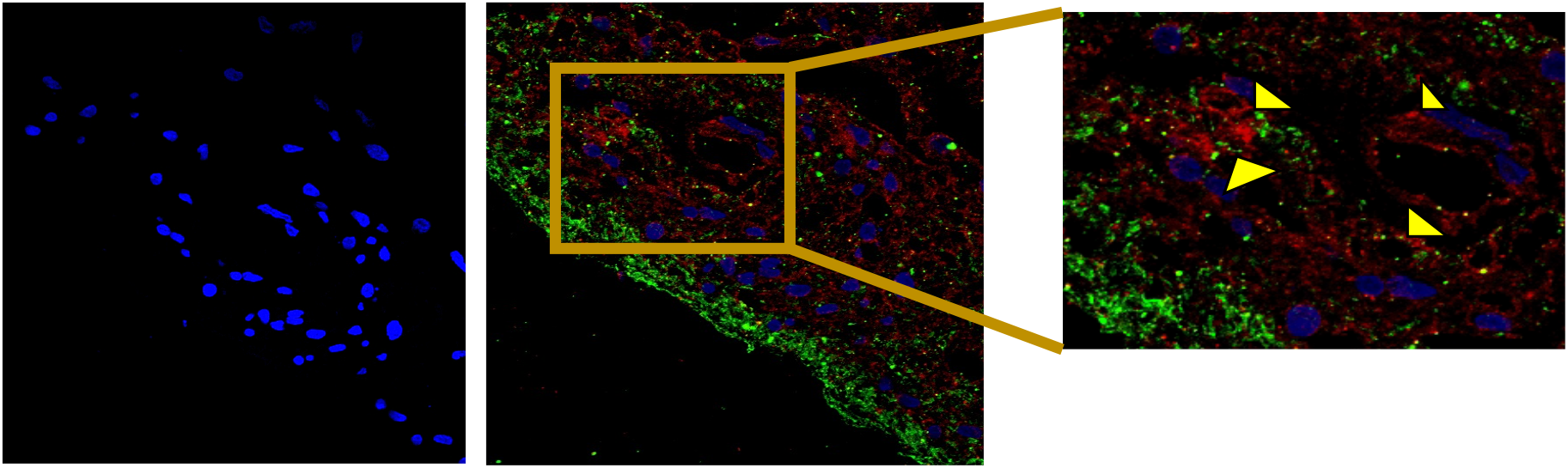
Immunohistochemistry of retinal section. (A) DAPI staining of the retinal section. (B) Merge image of EPHX2 and GFAP showing expression of both the proteins when stained with DAPI. (C) Enlarge image showing the expression of EPHX2 in and around retinal blood vessels when colocalized with GFAP.

**Figure 9:**
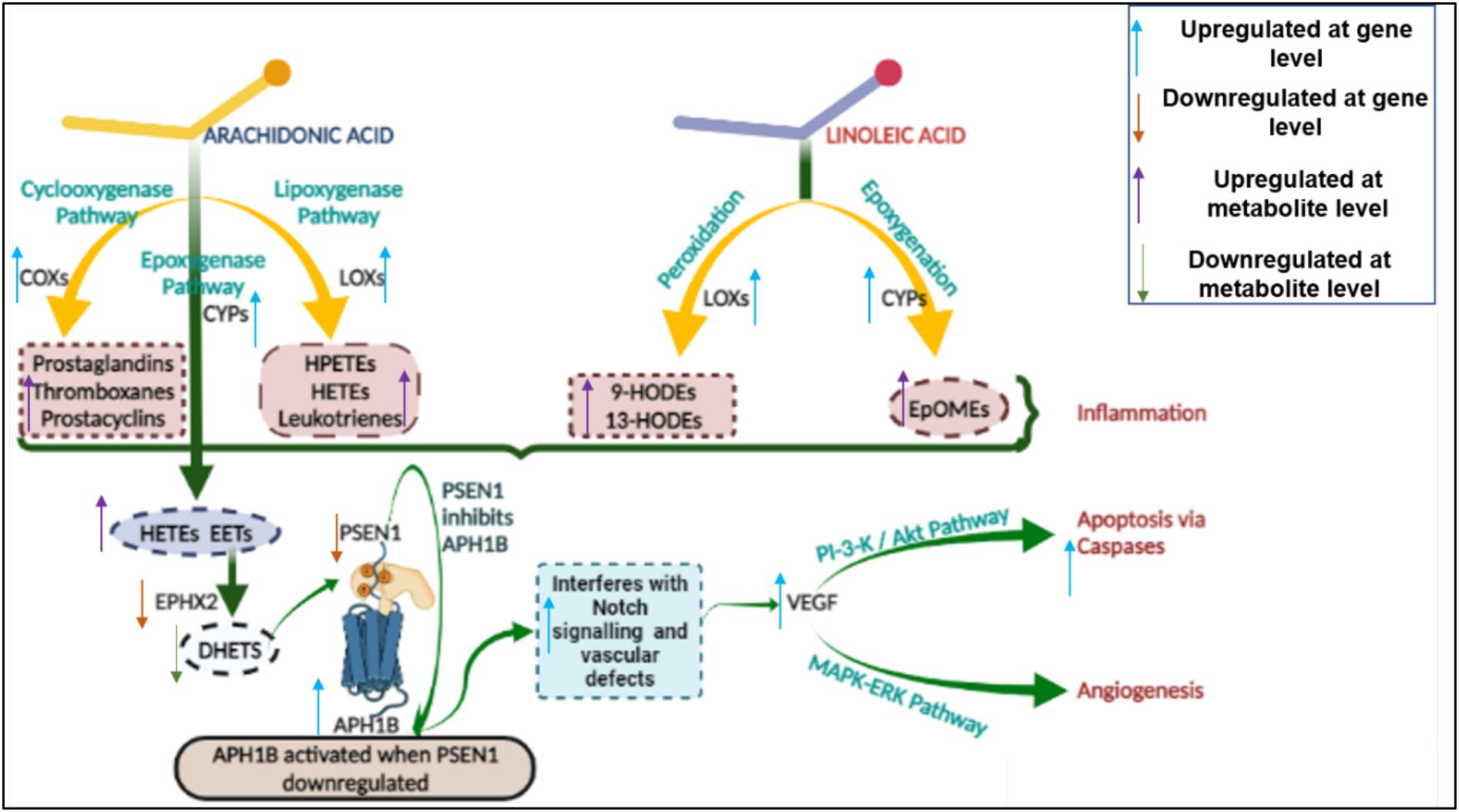
Schematic diagram of the proposed mechanism of altered lipid metabolism and its regulation by different proteins and enzymes. Upregulation of COXs, LOXs, and CYPs leads to accumulation of inflammatory lipid metabolites while EPHX2 downregulation leads to low dihydroxy fatty acid synthesis and further activating γ-secretase. Activation of notch receptor and VEGF enhances angiogenesis and apoptosis via the MAPK-ERK pathway and PI3K-Akt pathway respectively. The Blue and orange arrows denote upregulation and downregulation at mRNA expression levels respectively while the violet and green arrow denote upregulation and downregulation at vitreous metabolite expression levels respectively.

## Discussion

Several factors such as oxidative stress, inflammation, neovascularization, and apoptosis affect the overall development and maturation of retina in the preterm-born infants and contribute to ROP pathogenesis. These factors are known to be modulated by changes in lipid metabolism. PUFAs, a class of lipid, is abundantly present in the photoreceptor cells (rods and cones), macular layer, and nerve-fiber cells of the retina (15). Lipoxygenases (LOXs) such as ALOX15, ALOX5, and ALOX8, etc. are responsible for the stereo-specific peroxidation of PUFAs to their corresponding hydroperoxy derivatives (16). These LOXs are known to catalyze PUFAs by synthesizing 9, 12, 13-TriHOME, abundantly expressed by hematopoietic and immune cells, and thus are involved in inflammation (17). PUFAs are catalyzed by a group of Cytochrome P450 enzymes such as CYP1B1, CYP2C8, CYP2J11, and CYP2J9, etc. to yield epoxy fatty acids and ω-hydroxylated HETEs (6-, 17-, 18-, 19-, and 20-HETE). These epoxy fatty acids are catalyzed by soluble epoxide hydrolase (sEH) from diols fatty acids such as DHET (14, 15-dihydroxyeicosatrienoic acids), DHDP (19, 20-dihydroxydocosapentaenoic acid), and DHEQ (17, 18-dihydroxy-eicosatetraenoic acid), etc. which further down-regulate notch pathway. The significance of lipid-derived metabolites and the genes involved in their regulation in the pathogenesis of ROP has not been studied in detail. Thus, the major goal of the present study was to explore the role of lipid-derived metabolites and targeted enzymes/genes in contributing to increased inflammation and vascular changes as seen in ROP eyes. Hydrolase activity of soluble epoxide hydrolase is a major step in fatty acid metabolism that mediates inflammation as well as angiogenesis through Notch signaling (18, 19). Reduced expression /activity of sEH under hypoxia, affects DHDP and DHET (diols) conversion which further leads to loss of astrocytes in the retina that are required to guide and support growing vessels (10).

sEH (encoded by the EPHX2 gene), is one of the major steps in the AA and LA metabolism that is known to regulate Notch signaling, therefore we explored the role of the *EPHX2* gene and epoxy derived metabolites in ROP pathogenesis. An interaction map for the *EPHX2* gene showed its interactions with inflammatory mediated metabolites and CYP genes (CYP1B1, CYP2C8, etc.). These interactions are involved in the different pathways including arachidonic acid metabolism, Di-unsaturated fatty acid beta-oxidation, leukotriene metabolism, linoleate metabolism, and putative anti-inflammatory metabolites formation from EPA and xenobiotics metabolism pathways. Besides these metabolic interactions, the other interacting genes involved in angiogenesis *VEGF165, VEGF189, APH1B*, and *Notch1* were also significantly upregulated in preterm infants diagnosed with severe ROP while *PSEN1 was* significantly downregulated. VEGF165, a more potent regulator of angiogenesis and vasculogenesis (20) showed a higher fold change (>5) as compared to VEGF189.

*APH1B* (γ-secretase subunit gene) plays a role in angiogenesis by regulating Notch activity (21) while VEGF increases γ-secretase activity and Notch 1 cleavage. The inhibition of γ-secretase leads to decreased angiogenesis and blocks VEGF-induced endothelial cell proliferation, migration, and survival (22–24). A γ-secretase complex is composed of four different integral membrane proteins: presenilin1 (PS), nicastrin, Aph-1, and Pen-2. *PSEN1* (presinilin1) inhibits the cleavage of the APH1 domain that activates the transcription factor for Notch, thereby regulating angiogenesis (25). Downregulation of the PSEN1 gene could imply the cleavage of APH1B from the tetrameric complex, together with increased γ-secretase levels as regulated by VEGF further contributing to angiogenesis.

Notch is conventionally considered to be a controller of cell division, differentiation and pattern development, many studies have shown its role in developmental, adult, and pathological angiogenesis (26). It may be also noted here that of the Notch receptors, only Notch1 and Notch4 are expressed in vascular endothelial cells, which signifies a close association between Notch 1 and VEGF in angiogenesis. Diol fatty acids such as DHET and DHDP act on the presinilin1 domain of gamma-secretase to regulate the Notch signaling pathway (10, 27). Concurrent with a significantly higher expression of *CYP2C8, CYP1B1, COX2*, and *ALOX15* and lower expression of *EPHX2*, we found a high expression of metabolites generated from AA and LA (Prostaglandin, Thromboxane, DiHOMEs, TriHOMES, HODEs, HETEs, HEPEs, Epoxy fatty acids, etc.) and lower expression of *EPHX2* in ROP vitreous, suggesting further that a reduced expression of the diol (DHETs and DHDPs) affects angiogenesis through Notch pathway. In general, the Notch pathway is enhanced at low diol concentration and is reduced at high diol concentration. Thus, an overexpression of Notch could lead to abnormal angiogenesis or neovascularization as seen in the severe ROP stages.

Besides regulating angiogenesis, VEGF is also known to phosphorylate proteins involved in apoptosis through the Akt-PI3K pathway. The present study also found an upregulation of both initiator Caspase (*CASP3* gene) and executioner Caspase (*CASP8* gene), which further contributes to apoptosis. Earlier reports demonstrated that phosphatase activity of sEH leads to the production of nitric acid by phosphorylation of eNOS which regulates VEGF-mediated modulation in mammalian endothelial cells (28). This pathway may also induce apoptosis of neurons leading to higher inflammation as shown in different studies on ROP including the present study.

The biosynthesis of linoleic acid-derived oxylipins and trihydroxy-octadecenoic acids (TriHOMEs) is poorly understood in humans. sEH plays a significant role in the synthesis of TriHOMEs. TriHOMEs were found to be significantly upregulated (fold change= 4.15, p-value= 0.096) in ROP vitreous despite a reduced expression of sEH and this could possibly be due to the fact that TriHOME can either be generated both by enzymatic activity of sEH and/or non-enzymatic epoxy-alcohol formation by hydroperoxide isomerization as shown by Fuchs *et al*. (29). Likewise, while the epoxy metabolites EETs were observed in ROP vitreous, the absence of other epoxide fatty acids such EEQs and EDP could be explained as endogenous epoxide fatty acids are known to have a short half-life and highly unstable (30).

*EPHX2* genes are expressed by astrocytes and Muller glial cells that are known to guide angiogenesis by providing a scaffold for the endothelial cell to proliferate and form retinal blood vessels. We explored their expression in the glial cells present in the fibrovascular membrane (FVM), formed at the vitro-retinal interface in the eyes with tractional retinal detachment (TRD)/severe ROP. FVM is highly rich in lipid deposition and various cell types such as retinal glial cells, macrophages, monocytes, hyalocytes, laminocytes, fibroblasts, pericytes, and vascular endothelial cells (31, 32). *EPHX2* which is known to be expressed in astrocytes and Muller glial cells (10), did not show expression when co-localized with GFAP. However, expression of sEH was observed in the glial cells of the control retinal tissue, indicating the altered lower enzymatic expression in severe ROP infants.

Arachidonic acid and linoleic acid-derived metabolites are further categorized into inflammatory and anti-inflammatory-derived metabolites. Our study found the inflammatory metabolites from PUFAs, to be significantly upregulated (M335T27, M351T30, M299T27_1), while no significant change was seen in the anti-inflammatory fatty acids metabolites. HETEs generated from the activity of CYP1B1 and LOXs enzymes are known to activate neutrophils and macrophages and these were found to be upregulated in ROP infants. This further confirms the results from our previous study that demonstrated the presence of activated neutrophils and macrophages in ROP vitreous humor possibly leading to inflammation (Rathi et al. 2017).

The role of inflammation in the progression of ROP is well established. Inflammatory prostanoids such as prostaglandins, HETEs, and leukotrienes were upregulated while anti-inflammatory metabolites (DHETs and dihydroxy stearic acid) were downregulated in the ROP vitreous. Further correlation of the expression of angiogenesis-related and lipid metabolizing enzymes/genes, their metabolites with severe ROP, and their expression in the vitreous/FVM of severe ROP infants confirmed its possible role in mediating neovascularization. Further validation of the gene expression across different stages of ROP would have been ideal for studying a severity dependant change in lipid metabolizing enzymes and their activity, however, this was not possible due to the non-availability of vitreous samples from mild ROP. Astrocytes and other glial cells express epoxide hydrolase in the retina, thus performing a comparative expression of epoxide hydrolase across different cell types among cases and controls by immunostaining could have helped in distinguishing its function under disease vs normal state but this too could not be done due to non-availability of the retina from normal paediatric donors. Nevertheless, notwithstanding these limitations, the study still very clearly explained the involvement of fatty acid (PUFAs) metabolizing enzymes in ROP. This study can further be extended to study the response to ω-3 fatty acid supplementation in prenatal and early postnatal stages to mothers to identify the infants who may benefit from this treatment. Identifying and treating the disease at an early onset stage would certainly help in the appropriate and effective supervision of the disease and its progression. The findings of this study would aid in discovering lipid-based biomarkers for prognostic testing and detecting newer drug targets for efficient management of ROP.

## ETHICS STATEMENT

The study protocol adhered to the tenets of the declaration of Helsinki and written informed consent was obtained from the parents of all the minor subjects and was approved by the Institutional Review Board (LEC02-14-029) of the L V Prasad Eye Institute (LVPEI).

## AUTHOR CONTRIBUTIONS

IK and SK conceived the idea; IK, SJ, MBJ, and SC wrote the protocol; IK served as principal investigator; SC, SJ, MBJ, KA, and RK were co-investigators; SJ, RK, and KA performed clinical examinations, graded the fundus images and did surgeries for the preterm and full-term babies; SK and SP collected blood, vitreous and documented family history in the predesigned questionnaires; SK performed most of the molecular biology-based analysis of blood and membrane, MBJ, and SP performed mass spectrometry; SK, SP, IK, MBJ, and SC analyzed the data and wrote the manuscript; and all authors revised the paper and approved the submitted version.

## ACKNOWLEDGMENTS

The authors thank the parents of all the preterm and full-term babies for their voluntary participation. The authors would like to thank Mr. Sreedhar Rao Boyenpally for providing skin tissue and Ms. Suryadipali Pahadsingh for sample collection.

---

**Supplemtary table 1**: Primer sequences for the genes analysed for differential expression analysis by QPCR

