## Supplementary table 1 for "Detailed investigation on the role of lipid metabolizing enzymes in the pathogenesis of retinopathy of prematurity among preterm infants"

| Gene Name | Primer Sequence_Forward | Primer Sequence_Reverse |
|---|---|---|
| 1. *COX2* | 5’-CCTTCTGCCTGACACCTTTC-3’ | 5’-CAGGATACAGCTCCACAGCA-3’ |
| 2. *ALOX5* | 5’-AACGTGGAGCCTTCCTAACC-3’ | 5’-GGTGACAAAGTGGCAAACCT-3’ |
| 3. *CYP1B1* | 5’-CACGCCCCTTCTCTTCTCC-3’ | 5’-GACACCACACGGAAGGAGG-3’ |
| 4. *CYP2C8* | 5’-GAGACTGAGCCTAAGAACCCC-3’ | 5’-GAGGTCTTTTCTCCAGACTTGTT-3’ |
| 5. *EPHX2* | 5’-GGAAGTGTTATGTAAGGAGATG-3’ | 5’-GTAGAAGAGAGCCATGTACC-3’ |
| 6. *PSEN1* | 5’-GGTCCACTTCGTATGCTGGT-3’ | 5’-CGCAGGATACTGCGTGAAAG-3’ |
| 7. *APH1B* | 5’-GTGGGCATTCATGGAGATTC-3’ | 5’-GAAGCCTGAAACTCTGCCTG-3’ |
| 8. *VEGF165* | 5’-ATCTTCAAGCCATCCTGTGTGC-3’ | 5’-CAAGGCCCACAGGGATTTTC-3’ |
| 9. *VEGF189* | 5’-ATCTTCAAGCCATCCTGTGTGC-3’ | 5’-CACAGGGAACGCTCCAGGAC-3’ |
| 10. *NOTCH1* | 5’-TTGGGAGGAGCAGATTTTG-3’ | 5’-CACTGGCATGACACACAACA-3’ |
| 11. *CASP8* | 5’-GGAGCTGCTCTTCCGAATTA-3’ | 5’-GCAGGTTCATGTCATCATCC-3’ |
| 12. *CASP3* | 5'-ACATGGCGTGTCATAA AATACC-3' | 5'-CACAAAGCGACTGGATGAAC-3’ |
| 13. *β-ACTIN* | 5’-CATGTACGTTGCTATCCAGGC-3’ | 5’- CTCCTTAATGTCACGCACGAT-3’ |

Supplementary Table1: Sequence of forward and reverse Primers. List of the primer sequence for genes of lipid metabolism, angiogenesis and apoptosis. β-Actin was used as housekeeping gene.
